# Engineering pH-Responsive trans-Ferulic Acid/κ-Carrageenan Beads for On-Demand Micronutrient Delivery in Plants

**DOI:** 10.64898/2026.02.04.703679

**Authors:** Hitasha Vithalani, Subhojit Ghosh, Harshil Dave, Kashish Agrawal, Mukesh Dhanka, Subramanian Sankaranarayanan

## Abstract

Micronutrient deficiencies in soils are a critical challenge in agriculture, particularly in acidic soil environments where nutrient availability is strongly limited by fixation, leaching, and altered metal speciation. These constraints contribute to inefficient nutrient uptake and reduced crop yields. Conventional micronutrient supplementation methods are often inefficient, environmentally harmful, and unsustainable, underscoring the need for smarter delivery systems tailored to soil pH conditions. In this study, we developed biodegradable, pH-responsive microbeads from κ-carrageenan (κ-CG) and trans-ferulic acid (TFA) for targeted micronutrient release. The κ-CG–TFA microbeads were synthesized via an eco-friendly process and optimized for size, morphology, stability, and nutrient retention. Characterization confirmed the successful incorporation of functional groups, while swelling, degradation, and release studies demonstrated efficient delivery of essential micronutrients (Mn^2+^, Zn^2+^, Cu^2+^, and Fe^3+^) under acidic conditions (pH 4.0), mimicking acidic soil environments. The inherent antioxidant activity of TFA conferred strong radical-scavenging capacity, further enhancing its functionality. Soil water and plant growth assays revealed that the microbeads improved micronutrient availability, significantly increased chlorophyll content and leaf area, promoted vigorous seedling growth, and caused no phytotoxic effects. Collectively, these findings establish κ-CG–TFA microbeads as a promising, eco-friendly platform for sustainable micronutrient delivery and stress reduction, thereby improving crop productivity in agriculture.

## Introduction

Micronutrient deficiency in agricultural soils is a major barrier to sustainable crop productivity and nutritional security. Unlike macronutrients such as nitrogen (N), phosphorus (P), and potassium (K), which are required in large amounts, micronutrients including zinc (Zn), iron (Fe), manganese (Mn), copper (Cu), molybdenum (Mo), and boron (B) are required only in trace quantities but are indispensable for plant metabolic activity. They function as cofactors for enzymes, stabilizers of protein structures, and regulators of hormonal signaling and stress responses(1). Deficiencies in these elements disrupt photosynthesis, respiration, and reproductive processes, leading to symptoms such as chlorosis, necrosis, stunted growth, and reduced yields. (2). Deficiencies in these micronutrients have severe deleterious effects on plant health, leading to symptoms such as chlorosis, necrosis, stunted growth, and poor reproductive development.

Global incidences of micronutrient-deficient soils are a serious menace to sustainable agriculture and food security. About 30-50% of the earth’s arable land lacks zinc and iron, making these micronutrient deficiencies among the most prevalent in the world. Such deficiencies of soil nutrients have long-term effects not just on crop production and soil fertility but also on human health. The persistence of such deficiencies not only undermines crop productivity and soil fertility but also contributes to human malnutrition through the phenomenon of “hidden hunger,” in which caloric requirements are met but essential vitamins and minerals are lacking. Addressing this challenge has been identified as a priority by the Food and Agriculture Organization (FAO) and the World Health Organization (WHO) through strategies such as biofortification and improved nutrient management (3,4).

India’s varied agro-climatic conditions and intensive cultivation practices have resulted in widespread micronutrient deficiencies in soil, posing challenges to agricultural productivity and human nutrition. Diagnosis under the Soil Health Card scheme indicated that almost 39% of Indian soils are zinc-deficient, and approximately 19% are iron-deficient (4). Variations across states are significant, with states such as Bihar, Odisha, and Gujarat among the highest for micronutrient-deficient soils. Factors such as monocropping, unbalanced NPK-dominated fertilizer use, limited application of organic manures, and residue removal further intensify these deficiencies. Moreover, alkaline, calcareous, and salt-affected soils typical of arid regions exacerbate micronutrient immobilization, reducing bioavailability (5).

The rhizosphere, the narrow soil zone surrounding plant roots, plays a central role in nutrient acquisition. Plants actively modify this environment through the secretion of root exudates, including organic acids, amino acids, sugars, flavonoids, and enzymes, which modulate pH, solubilize bound nutrients, and recruit beneficial microorganisms (6). One of the primary adaptive mechanisms in micronutrient deficiency is acidification of the rhizosphere by proton extrusion and secretion of organic acids. This reduces the local pH, enhancing the solubility of normally immobile micronutrients such as Fe^3+^, Zn^2+^, and Mn^2+^ (7). However, prolonged acidification in nutrient-poor soils can adversely affect soil microbial communities and degrade long-term fertility (8). Microbes associated with the rhizosphere are crucial to micronutrient mobilization. Certain soil microbes produce siderophores, low-molecular-weight compounds with a high affinity for iron, solubilizing it for plant uptake. Others secrete organic acids and phosphatases that liberate micronutrients from mineral-bound forms (9). While biofertilizers and microbial inoculants provide a natural approach to enhancing nutrient availability, their performance in the field is often constrained by soil health, competition with indigenous microbiota, and environmental variability.

Conventional approaches to correcting soil micronutrient deficiencies include soil application of inorganic salts, foliar sprays, chelated formulations, and coated fertilizers. These methods, however, face significant limitations: precipitation of nutrients into insoluble forms in alkaline soils (10). Foliar sprays give instant correction but are not feasible for large fields and induce phytotoxicity. Chelated micronutrients, while effective, are costly and environmentally persistent (11). Micronutrient-coated fertilizers are generally plagued with irregular release rates and low coating efficiency. New technologies, such as microbial biofertilizers and nano-fertilizers are still mostly experimental and suffer from scalability, regulatory, and ecological risk assessment challenges (12).

To overcome these constraints, “smart” nutrient delivery systems that respond to environmental triggers have gained attention. Such systems include materials and formulations designed to be sensitive to specific environmental cues, such as pH, temperature, or moisture levels, to release nutrients only when and where they are required. Of these, pH-sensitive systems are especially appropriate for the delivery of micronutrients in soils, as root exudates actively lower rhizosphere pH under nutrient stress. Carriers that can sense acidification and release encapsulated micronutrients provide a highly targeted and effective mechanism for nutrient delivery (13,14). Carriers that respond to this localized pH reduction offer a highly targeted mechanism for micronutrient release, improving nutrient use efficiency (NUE), reducing leaching losses, and minimizing environmental impacts such as eutrophication and soil degradation.

In this study, we report the development of a biodegradable, pH-sensitive nutrient delivery system fabricated from two natural and biocompatible materials: trans-ferulic acid (TFA) and κ-carrageenan (κ-CG). TFA, a hydroxycinnamic acid widely distributed in cereals such as oats, rice, and wheat, was selected as the functional crosslinker and bioactive additive (15). Its ionizable phenolic and carboxylic groups impart pH-responsiveness, while its capacity to form hydrogen and ester bonds with polysaccharides enhances the mechanical stability of hydrogel matrices (16). Additionally, TFA possesses intrinsic antioxidant activity, shielding encapsulated micronutrients from oxidative degradation during storage and application, thereby preserving bioavailability (17). Being eco-friendly and biodegradable, TFA degrades without leaving toxic residues, making it suitable for sustainable agricultural applications (18). Its pH-sensitive ionizable phenolic and carboxylic groups make it suitable for pH-responsive systems. κ-Carrageenan (κ-CG), a sulfated seaweed polysaccharide, is utilized across food and biomedical industries for its stabilizing and gel-forming capabilities (19,20). Agriculturally, κ-CG is valued for its pH-sensitive sol-gel transitions, high ion-binding capacity, biodegradability, and compatibility with polyphenols. Its sulfate moieties strongly interact with metal cations such as Fe^3+^, Zn^2+^, Mn^2+^, and Cu^2+^, facilitating efficient encapsulation and controlled delivery (21). In blends with TFA, κ-CG generates interpenetrating polymer networks (IPNs), increasing the structural integrity, toughness, and shelf-life of microbeads (22).

In this study, we developed a biodegradable, pH-responsive nutrient delivery system (**Scheme 1**) using trans-ferulic acid (TFA) and κ-carrageenan (κ-CG), selected for their natural origin, biocompatibility, and complementary functions. The κ-CG-TFA microbeads were engineered to exploit rhizospheric acidity, enabling controlled release of essential micronutrients where deficiencies are most severe. This strategy not only enhances nutrient bioavailability and plant uptake but also reduces leaching losses, offering a sustainable route to improved crop productivity and soil health. Importantly, the system integrates structural stability, antioxidant functionality, and environmentally triggered nutrient release in a single platform, thereby aligning with the principles of smart and sustainable agriculture. Collectively, this work introduces a next-generation fertilizer concept with potential to mitigate micronutrient deficiencies and strengthen agricultural resilience.

**Scheme 1..**
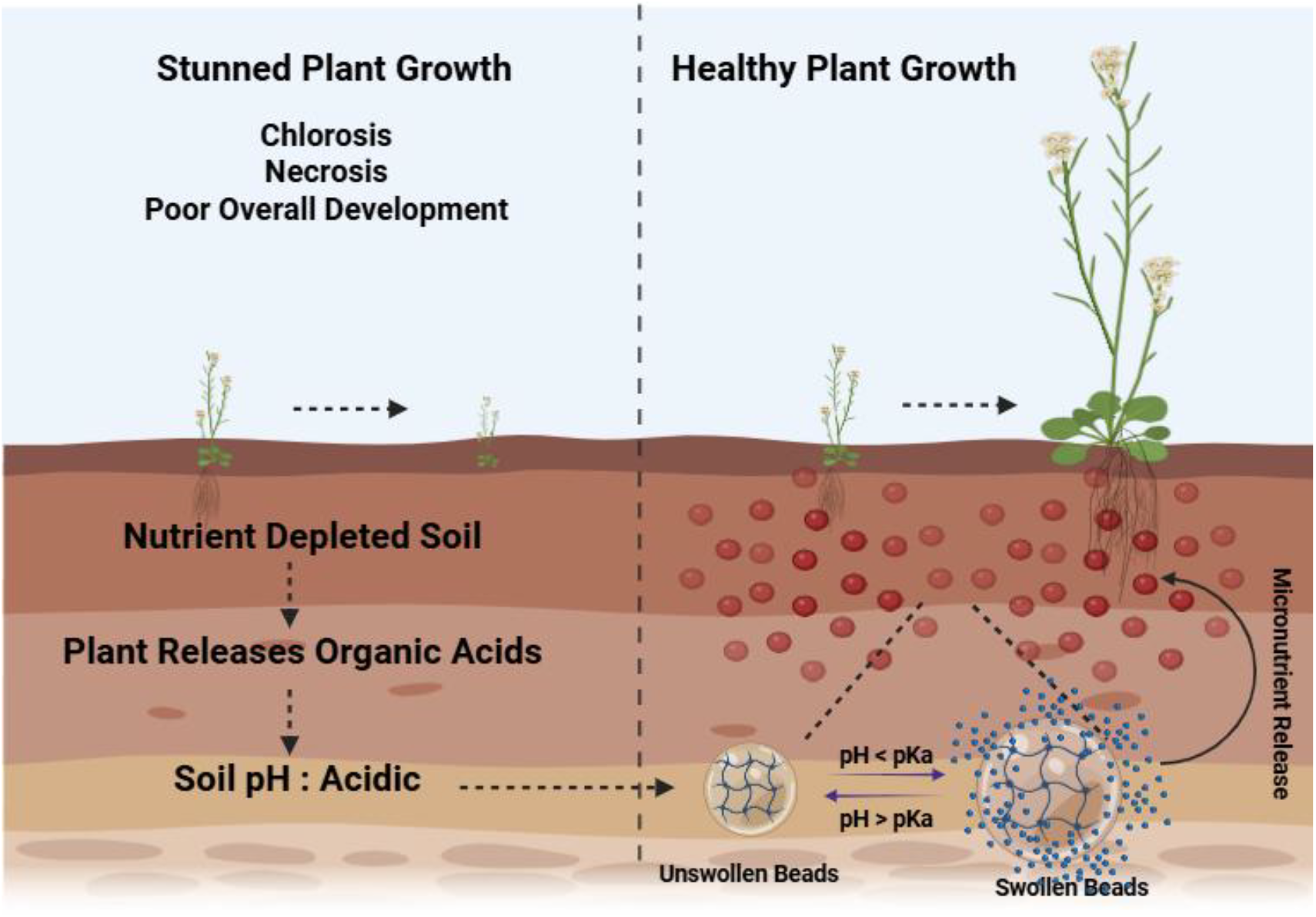
Schematic illustration of the κ-carrageenan-trans-ferulic acid microbead system for pH-responsive micronutrient delivery in soil. In nutrient-depleted and acidic soils, plants release organic acids that lower rhizospheric pH. The microbeads swell under acidic conditions (pH < pKa), triggering controlled release of encapsulated micronutrients (Mn^2+^, Zn^2+^, Cu^2+^, Fe^3+^), which alleviate chlorosis, necrosis, and stunted growth, thereby restoring healthy plant development.

## 2. Experimental Section

### 2.1 Materials

Trans-ferulic acid (TFA) and κ-carrageenan (κ-CG) were obtained from TCI, India. Metal salts used for micronutrient encapsulation included manganese chloride (MnCl_2_), zinc nitrate hexahydrate [Zn(NO_3_)_2_·6H_2_O], cupric sulphate (CuSO_4_), and ferric chloride (FeCl_3_), sourced from Sigma-Aldrich and SRL, India. Calcium chloride (CaCl_2_·2H_2_O) was used as a crosslinking agent. All chemicals were of analytical grade. Deionized water was used throughout. MS (Murashige and Skoog 1962) medium and MS macronutrients have been acquired from Himedia Laboratories. All other reagents used in this study were of analytical grade or above.

### 2.2 Synthesis of TFA-κ-CG Microbeads and Encapsulation of micronutrients

Microbeads were prepared by a modified ionotropic gelation technique. κ**-**Carrageenan (2% w/v) was dissolved in hot distilled water (90 °C) under continuous stirring until a clear and homogeneous solution was obtained. trans-Ferulic acid (0.05% w/v), pre-solubilized in minimal warm water, was added to the polymeric solution and allowed to integrate fully into the carrageenan matrix. Subsequently, micronutrient salts (MnSO_4_, ZnSO_4_, CuSO_4_, and FeSO_4_) were individually incorporated into the TFA-κ-CG solution to obtain four formulations, designated as K-TF-Mn, K-TF-Zn, K-TF-Cu, and K-TF-Fe. The final mixture was transferred into a syringe and extruded dropwise into a chilled crosslinking bath containing 10% (w/v) CaCl_2_ supplemented with the respective micronutrient salt. Upon contact, spherical beads formed instantly due to ionic crosslinking between κ-carrageenan sulfate groups, calcium ions, and micronutrient cations. The beads were allowed to harden in the crosslinking bath for 1-2 h to ensure structural stability, collected by filtration, washed with distilled water to remove unbound ions, and dried at 40 °C for 24 h. Blank control beads (without micronutrients) were prepared following the same procedure.

### 2.3 Physicochemical Characterization

To confirm bead formation and elucidate the functional groups involved in crosslinking and micronutrient encapsulation, Fourier Transform Infrared (FTIR) spectroscopy was carried out using a Shimadzu FTIR-8400S spectrometer. The characteristic absorption bands were analyzed to verify the presence of κ-carrageenan and trans-ferulic acid, as well as their interactions with incorporated micronutrients within the bead matrix.

The internal structure and cross-sectional morphology of the beads were further examined by scanning electron microscopy (SEM), providing insight into the network’s compactness and the distribution of the polymeric matrix after crosslinking.

Bead size was determined for all formulations using an inverted optical microscope (Leica DMi8), and the same system was employed to evaluate the overall morphology and surface texture of the synthesized microbeads, including shape uniformity, surface smoothness, and structural integrity.

### 2.4 Swelling and Degradation Studies

The pH-responsive swelling behavior of the microbeads was evaluated by immersing 100 mg of each dried formulation in 10 mL of buffer solutions at pH 4.0, 7.0, and 9.0. Samples were incubated at 25 °C for 24 hours. At predetermined time intervals (0, 1, 2, 4, 8, 12, and 24 h), beads were retrieved, gently blotted to remove surface moisture, and weighed. The swelling ratio (SR) was calculated using (23):

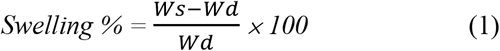

Where, Ws is the weight of swollen beads and Wd is the weight of dried beads.

For degradation studies, 100 mg of beads were immersed in the same buffer systems and incubated for 1-14 days. At defined time points, beads were collected, washed, dried to constant weight, and weighed. The percentage degradation (%) was calculated as (23,24):

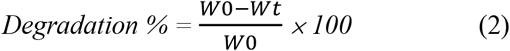

where, W0 is the initial weight of microbeads and Wt is the remaining weight of microbeads after degradation.

### 2.5 pH-Triggered Release Behavior of Phenol Red Encapsulated Beads

The pH-responsive release of phenol red from κ-carrageenan/trans-ferulic acid beads was evaluated in buffer solutions at pH 4.0 (acidic), 7.0 (neutral), and 9.0 (basic). A fixed mass of dye-loaded beads was incubated in the respective release media at room temperature under gentle shaking. The study was conducted for 21 days, with samples collected at daily intervals. At each time point, a defined volume of the medium was withdrawn and replaced with fresh buffer to maintain sink conditions. The released phenol red was quantified using a UV-visible spectrophotometer (GENESYS 50) at 430 nm for pH 4 and 560 nm for pH 7 and pH 9. Dye concentration was determined from pH-specific calibration curves. In parallel, bead size was measured at each pH condition using optical microscopy to correlate swelling behavior with release profiles. Cumulative release profiles were calculated as a function of time. All experiments were performed in triplicate. The cumulative release percentage at each time point was calculated as (23,24) :

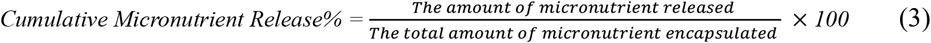

### 2.6 Antioxidant Activity

The antioxidant potential of the microbeads was assessed using the 2,2-diphenyl-1-picrylhydrazyl (DPPH) radical-scavenging assay. A 200 μM DPPH solution was freshly prepared in methanol. To obtain the test samples, the beads were immersed in distilled water overnight, allowing the release of encapsulated micronutrients and trans-ferulic acid into the leachate. Aliquots of the leachate were then mixed with an equal volume of the DPPH solution and incubated in the dark at room temperature for 30 minutes. The decrease in absorbance was recorded at 517 nm using a UV-Vis spectrophotometer (GENESYS 50).The percentage radical scavenging activity was calculated using the following equation:(23).

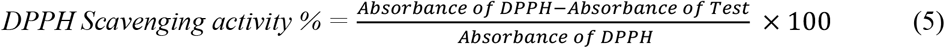

### 2.7 Plant Study

Plant growth conditions: *Arabidopsis thaliana* (Col-0 ecotype) and *Nicotiana benthamiana* plants were grown in a controlled growth chamber/growth room at 22 °C under a 16 h light/8 h dark photoperiod with 70-80 % relative humidity. Plants were cultivated in a soil mixture composed of soilrite, vermiculite, and perlite in a 3:1:1 ratio, and the soil pH was adjusted to 4.0 using HCl.

#### Measurement of leaf area

Leaf area was measured using Fiji (ImageJ, NIH, USA). Digital images of leaves were taken under uniform light conditions, and the scale was calibrated in Fiji using a reference measurement. The leaf outlines were then selected, and the software automatically calculated the leaf area.

#### Measurement of chlorophyll content

Chlorophyll is extracted from fresh plant tissue using 80 % acetone, as it is readily soluble in organic solvents. About 15 mg of finely cut leaf tissue is ground with 0.5 mL of 80% acetone in a centrifuge tube with a micro-pestle. The homogenate is centrifuged at 5,000 rpm for 5 minutes, and the supernatant is collected. The residue is re-extracted with 80 % acetone until it becomes colorless, and all extracts are pooled in a 2 mL volume centrifuge tube. The final volume is adjusted to 2 ml with 80 % acetone. Absorbance of the extract is recorded at 645 nm and 663 nm (with 80 % acetone as blank) using a spectrophotometer. The concentrations of total chlorophyll (mg per g fresh weight tissue) are calculated using the following equations (24):

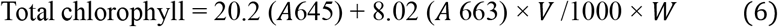

where A = absorbance at respective wavelengths, V = final volume of extract (mL), and W = fresh weight of tissue (g).

### 2.8 Statistical Analysis

All statistical evaluations were carried out using OriginPro 2017 (OriginLab, USA) with one-way ANOVA to assess differences among groups. Post-hoc comparisons were conducted using the Tukey-Kramer multiple comparison test. Data are expressed as mean ± standard deviation (SD), and most experiments were performed in triplicate. For in vivo studies, statistical analyses were performed with GraphPad Prism version 9 (GraphPad Software, USA) using one-way ANOVA followed by Tukey-Kramer post-hoc testing. Statistical significance levels were indicated as: *p < 0.05, **p < 0.01, ***p < 0.001, and ****p < 0.0001.

## 3. Results and Discussion

### 3.1 Synthesis of Microbeads and Encapsulation of Micronutrients

A systematic optimization was carried out by varying the concentrations of κ-carrageenan (κ-CG) and trans-ferulic acid (TFA) to achieve mechanically robust and pH-responsive microbeads. Ionotropic gelation efficiency was found to be highly dependent on the balance between the polysaccharide matrix and the phenolic additive as depicted in **Figure 1a**. At low κ-CG concentrations (<2% w/v), insufficient polymer chain density led to weak electrostatic crosslinking with Ca^2+^ ions, yielding fragile, irregularly shaped beads prone to deformation. Conversely, increasing the κ-CG concentration beyond 2% produced highly viscous solutions that hindered uniform droplet extrusion and led to non-spherical aggregates.

**Figure 1:**
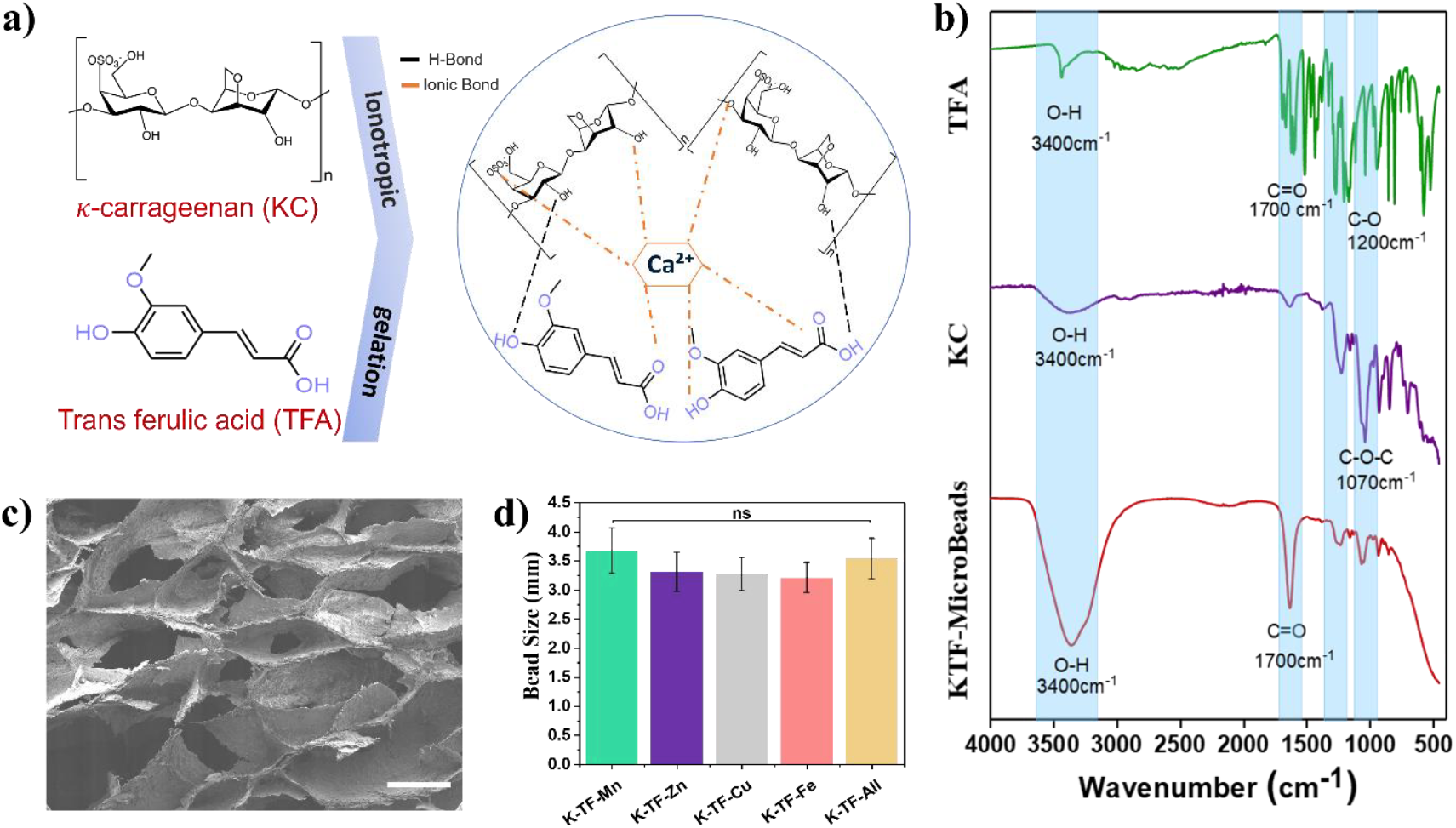
Schematic illustration depicting (a) the chemical structure of κ-carrageenan and trans-ferulic acid, the ionotropic gelation-based fabrication of composite beads, and the possible intermolecular interactions within the hydrogel network. (b) FTIR spectra components within the composite beads. (c) FESEM micrograph showing the surface morphology of the hydrogel beads (Scale bar: 100 μm). (d) Distribution of encapsulated micronutrients, indicating non-significant variation in bead diameter.

Similarly, the TFA concentration strongly influenced bead integrity. While incorporation of 0.05% TFA synergistically enhanced the structural stability through hydrogen bonding and π-π interactions with κ-CG chains, higher TFA levels (>0.1%) interfered with ionic crosslinking. This disruption likely arises from the competitive chelation and phenolic-calcium interactions, which weaken the sulfate-Ca^2+^ crosslinking necessary for bead hardening.

The optimized formulation (2% κ-CG with 0.05% TFA) produced spherical, smooth-surfaced beads with consistent size distribution and high resilience under mechanical handling. Importantly, this composition provided a balance between matrix rigidity (for stability in soil conditions) and environmental responsiveness (for pH-triggered release of micronutrients). Thus, the selected formulation ensured reproducibility, scalability, and functional performance for sustainable nutrient delivery applications.

Encapsulation of essential micronutrients (Mn, Zn, Cu, Fe) into the optimized κ-carrageenan/trans-ferulic acid (K-TF) matrix resulted in the formation of discrete, spherical microbeads with smooth surfaces and distinct ion-dependent colorations (**Figure 1d and 2a**). The beads exhibited average diameters between 3.21-3.68 mm, reflecting excellent control over droplet size and crosslinking kinetics. Morphological assessment revealed that K-TF-Zn beads demonstrated the highest uniformity and surface integrity, suggesting robust coordination between Zn^2+^ ions and the sulfate moieties of κ-carrageenan, stabilized further by the phenolic crosslinking of TFA. This synergistic ionic-phenolic interaction minimized bead deformation during gelation, yielding consistently monodisperse structures. In contrast, beads containing Mn^2+^, Cu^2+^, and Fe^2+^, although spherical, exhibited subtle variations in translucency and surface compactness, indicating differences in ionic affinity and gelation dynamics. Notably, multi-nutrient formulations (K-TF-All) displayed slightly irregular morphologies and broader size distribution, attributable to competitive ion binding within the carrageenan network. This observation underscores the role of ionic specificity in dictating bead integrity and highlights potential trade-offs in multi-micronutrient encapsulation systems. Collectively, these findings confirm that Zn-loaded K-TF beads provide an optimal balance between morphological stability, ion-polymer affinity, and encapsulation homogeneity, establishing them as a model system for subsequent nutrient release and soil-interaction studies.

**Figure 2:**
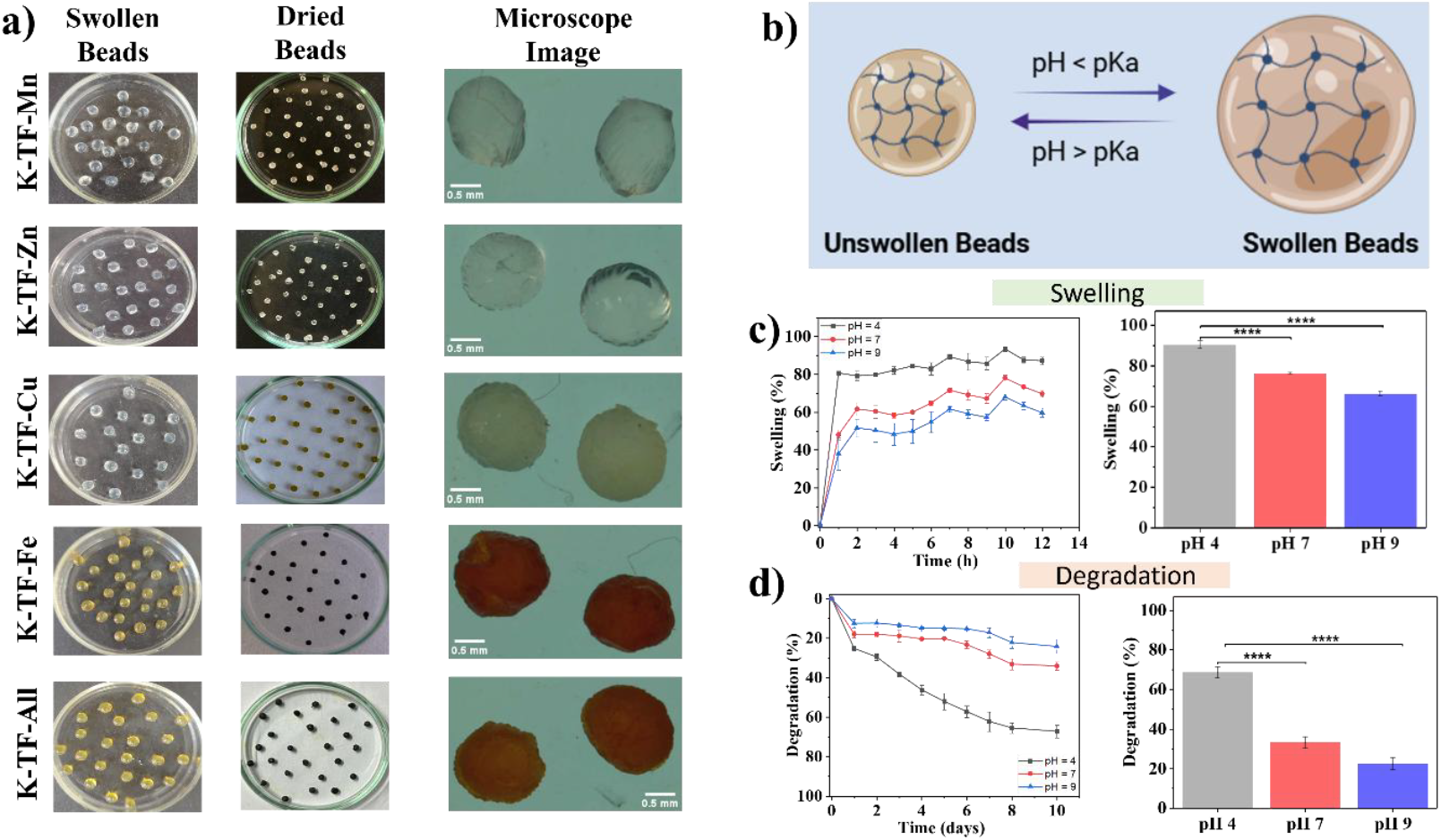
Morphological and physicochemical characterization of the hydrogel beads, showing (a) macroscopic and microscopic images of swollen and dried beads. (b) Proposed mechanism of bead swelling in an aqueous environment. (c) Swelling kinetics of the beads over time. (d) Degradation profile of the hydrogel beads at different pH.

### 3.2 Physicochemical Characterization

The FTIR spectra provide clear evidence of interactions between κ-carrageenan (κ-CG), trans-ferulic acid (TFA), and encapsulated micronutrients. Pure κ-CG (**Figure 1b**) exhibited its characteristic functional group vibrations, including a broad band at ∼3400 cm^−1^ (-OH stretching of hydroxyl groups), a sharp band at ∼1220 cm^−1^ (asymmetric stretching of sulfate esters, -SO_3_^−^), and an intense peak at ∼1070 cm^−1^ (C-O-C stretching of glycosidic linkages). These peaks serve as the spectral fingerprint of the polysaccharide backbone.

Upon incorporation of TFA (**Figure 1b**), new bands emerged in the region of ∼1600-1700 cm^−1^, attributable to aromatic C=C stretching and carbonyl (C=O) vibrations of the phenolic acid. The appearance and enhancement of these bands indicate successful integration of TFA within the κ-CG matrix through hydrogen bonding and electrostatic interactions, without disrupting the native polysaccharide framework.

When micronutrient salts were encapsulated (**Figure S1 a-d**), distinct spectral modifications were observed. The sulfate band (∼1220 cm^−1^) and carboxyl-related vibrations (∼1400-1600 cm^−1^) displayed shifts in peak position and relative intensity, consistent with ionic coordination between divalent cations (Zn^2+^, Mn^2+^, Cu^2+^, Fe^2+^, or multi-nutrient mixtures) and the negatively charged sulfate/carboxyl groups of κ-CG and TFA. These changes confirm the role of electrostatic crosslinking in bead stabilization and micronutrient entrapment.

Notably, all micronutrient-loaded beads (**Figure S1 a-d**) showed broadened -OH stretching bands, suggesting additional hydrogen bonding between hydroxyl groups of κ-CG/TFA and hydrated metal ions. Such interactions not only reinforce the structural integrity of the bead matrix but also contribute to the controlled release properties under acidic pH conditions.

As observed in **Figure 1c**, the SEM analysis confirms Ca^2+^-mediated crosslinking and hydrogen bonding between κ-carrageenan and trans-ferulic acid, cross-sectional SEM analysis revealed a well-defined porous internal structure. The beads exhibited interconnected pores separated by thin polymer walls, characteristic of ionotropically gelled κ-carrageenan networks. This sponge-like morphology provides diffusion pathways for encapsulated micronutrients. These spectral features support the proposed role of such interactions in stabilizing the bead structure and in enabling the observed pH-responsive release behavior.

### 3.3 pH-Responsive Swelling and Degradation

Swelling experiments (**Figure 2b-c**) demonstrated that all κ-CG-TFA microbead formulations displayed a distinct pH-dependent response. Maximum swelling of approx. 85 % occurred at pH 4, where the beads absorbed significantly more water than in neutral (73 %) or alkaline conditions (62 %). This behavior is attributed to enhanced ionization of carboxyl and hydroxyl groups under acidic conditions, which increases electrostatic repulsion within the hydrogel matrix and drives network expansion. Such targeted swelling is particularly advantageous for agronomic applications, since rhizosphere soils often exhibit weakly acidic microenvironments. By responding specifically to this niche, the beads ensure that nutrient release is spatially confined to the crop root zone.

Degradation analysis was recorded after the beads had reached saturated swelling. **Figure 2d** clearly demonstrated the pH-responsive degradation behaviour of the bead system. Maximum degradation was observed under acidic conditions, with approximately 65% degradation occurring at pH 4.0, indicating accelerated matrix breakdown. In comparison, lower degradation levels were recorded at neutral pH (∼33%) and under alkaline conditions (∼24%), reflecting greater structural stability of the beads. The enhanced degradation at acidic pH can be attributed to proton-induced weakening of Ca^2+^-κ-carrageenan ionic junctions and increased polymer chain mobility. This pH-dependent degradation trend is consistent with the swelling and release profiles, confirming the suitability of the system for controlled, on-demand micronutrient delivery.

Overall, the swelling and degradation results (**Figure 2b-d**) confirm that κ-CG-TFA beads act as a rhizosphere-adaptive, pH-responsive nutrient delivery system capable of providing sustained micronutrient release under acidic soil conditions.

### 3.4 pH-Responsive Release

The release kinetics of phenol red from κ-carrageenan-trans-ferulic acid (κ-CG-TFA) microbeads demonstrated pronounced pH dependence, underscoring their rhizosphere-adaptive functionality (**Figure 3a-d**). Phenol red was employed as a model dye to simulate real-time micronutrient release behavior. The schematic in **Figure 3a** illustrates the pH-responsive release mechanism. Under acidic conditions (pH 4.0), the microbeads exhibited a rapid and sustained increase in cumulative release over the 18-day period, reaching a maximum release of ∼85%, significantly higher than that observed in neutral and alkaline media. In comparison, a moderate release profile was recorded at pH 7.0 (∼47%), while the slowest and lowest release occurred at pH 9.0 (∼38%), indicating enhanced matrix stability under alkaline conditions (**Figure 3b-d**).

**Figure 3:**
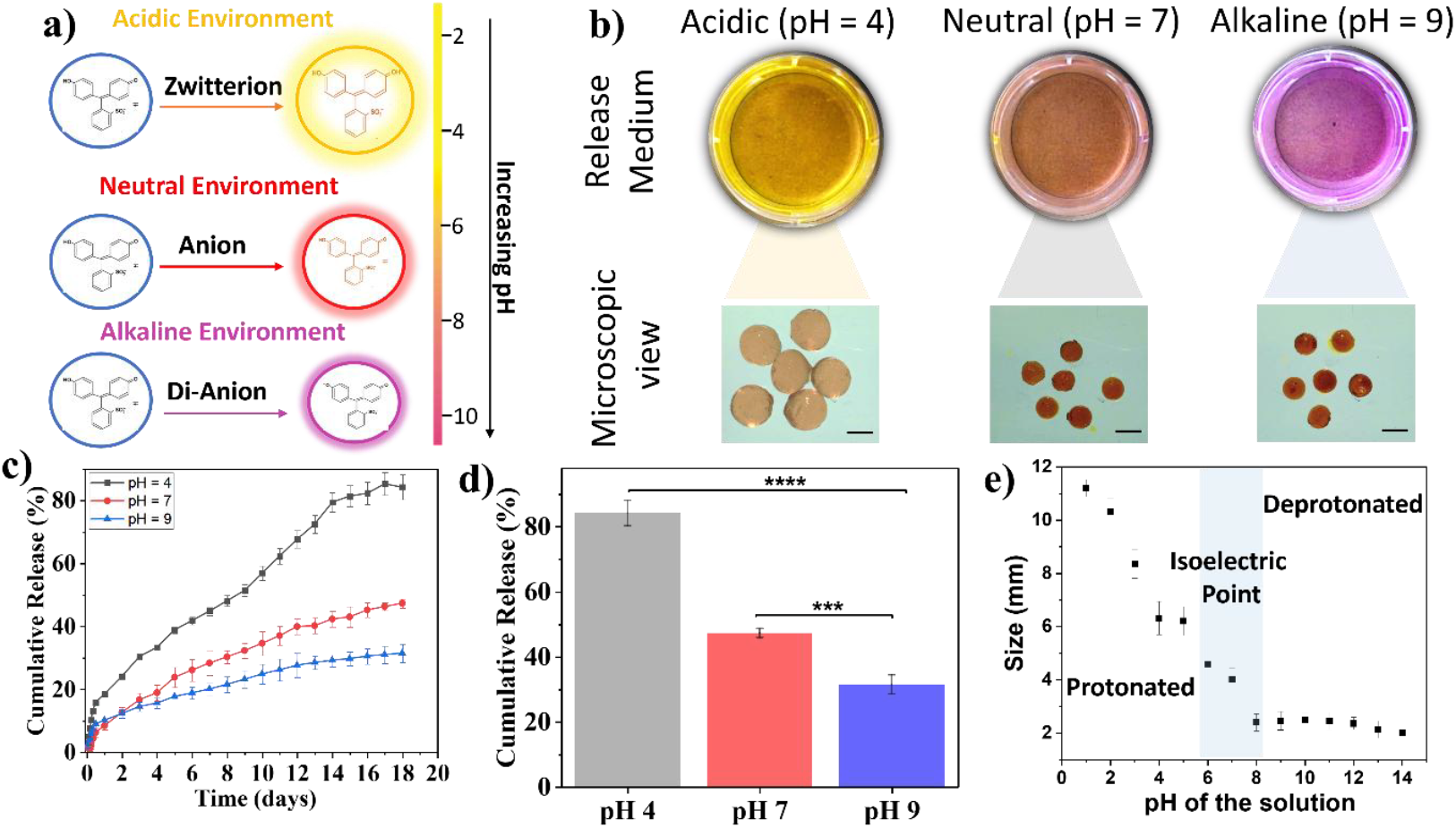
pH-responsive behavior and release characteristics of the hydrogel beads, where (a) a schematic representation illustrates the pH-dependent color change of phenol red and the associated release mechanism. (b) Cumulative release profiles in media of different pH levels, along with microscopic images of beads under corresponding pH conditions (Scalebar: 200µm). (c) Comparative release kinetics at different pH values. (d) Statistical analysis highlighting significant differences in release behavior after 21 days. (e) Variation in bead diameter as a function of pH.

This pH-dependent release behavior closely correlated with bead size variations measured under different pH environments. As shown in **Figure 3e**, optical microscopy revealed pronounced bead swelling under acidic conditions, with the largest bead diameter (∼11.5 mm) observed at pH 1.0. In contrast, reduced swelling was evident at pH 7.0, where the bead diameter decreased to ∼4.5 mm, and minimal swelling occurred at pH 9.0 and above, yielding the smallest bead size of ∼2.5 mm. The enhanced swelling under acidic conditions can be attributed to proton-induced weakening of Ca^2+^-κ-carrageenan-TFA ionic crosslinks, along with increased hydration of the polymer network, which promotes matrix relaxation and diffusion-controlled release. Conversely, stronger ionic interactions and restricted polymer chain mobility under neutral and alkaline conditions limited bead expansion and retarded dye diffusion.

Overall, the coupled swelling and release profiles confirm that κ-CG-TFA microbeads exhibit selective pH responsiveness, enabling controlled and on-demand micronutrient delivery, particularly suited for acidic rhizosphere environments.

### 3.5 Antioxidant Activity

The antioxidant potential of the microbeads was evaluated using the DPPH radical scavenging assay, with the underlying mechanism illustrated in **Figure 4a**. All formulations demonstrated substantial antioxidant activity, with K-TF-Zn (∼89%) and K-TF-Mn (∼85%) exhibiting the highest radical scavenging efficiencies, followed by K-TF-Cu (∼78%) and K-TF-Fe (∼77%). The pronounced antioxidant activity is primarily attributed to the phenolic structure of trans-ferulic acid (TFA), in which hydroxyl and methoxy functional groups promote effective electron donation, thereby neutralizing DPPH radicals. Notably, the blank microbeads also displayed considerable scavenging activity (∼75%), as shown in **Figure 4b-c**, indicating an inherent antioxidant contribution from the κ-carrageenan matrix itself, likely due to residual hydroxyl functionalities.

**Figure 4.**
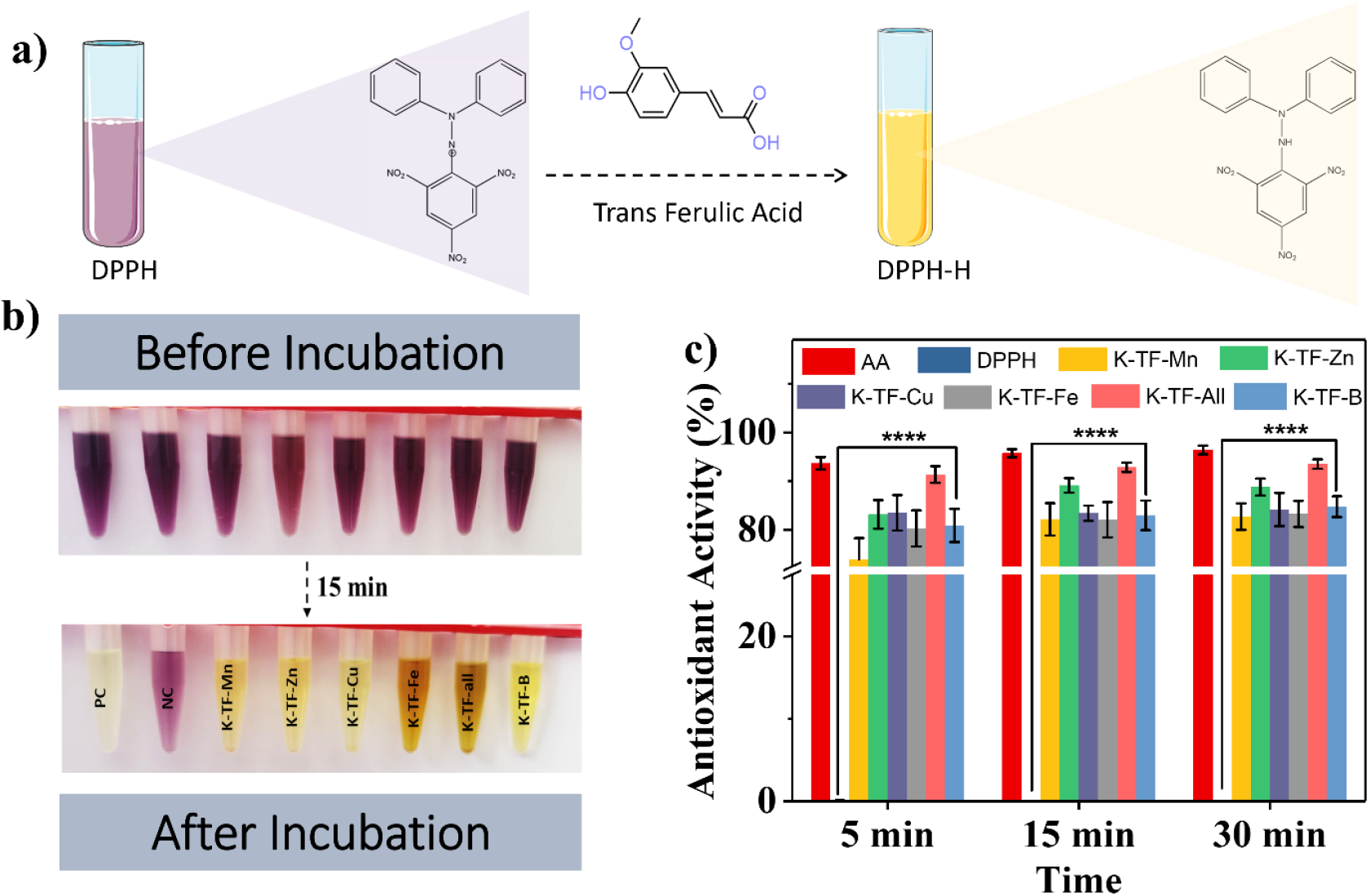
Antioxidant performance of the hydrogel beads demonstrated through (a) schematic illustration of the DPPH radical scavenging mechanism. (b) Representative photographs showing visual changes during the DPPH assay. (c) Quantitative graphical representation of DPPH radical scavenging activity. PC = positive control (ascorbic acid), NC = negative control.

The enhanced performance of K-TF-Zn is likely due to synergistic redox interactions between Zn^2+^ ions and phenolic groups, which amplify radical quenching. Conversely, the reduced activity of K-TF-Fe can be explained by the tight coordination of Fe^3+^ with phenolic hydroxyls, limiting their accessibility for radical neutralization. This trade-off suggests that Fe confers stability to encapsulated TFA but at the expense of antioxidant efficiency.

Overall, incorporation of TFA into κ-carrageenan microbeads imparts structural stability and intrinsic antioxidant functionality, thereby protecting micronutrients from oxidative degradation and potentially extending their shelf-life.

### 3.6 Effects on Plant Growth

All formulations were evaluated using two plant species, *Arabidopsis thaliana* and *Nicotiana benthamiana*, with the experimental methodologies illustrated in **Figure 5a** and **Figure 6a**, respectively. Plants were grown in soil with added nutrients. Positive control containing ½ MS complete media, negative control containing ½ MS macronutrients only. Other nutrient combinations are added with microbeads. Plant growth performance was assessed using morphological observations, leaf area measurements, and chlorophyll content as standard parameters for analysis and comparative evaluation.

**Figure 5.**
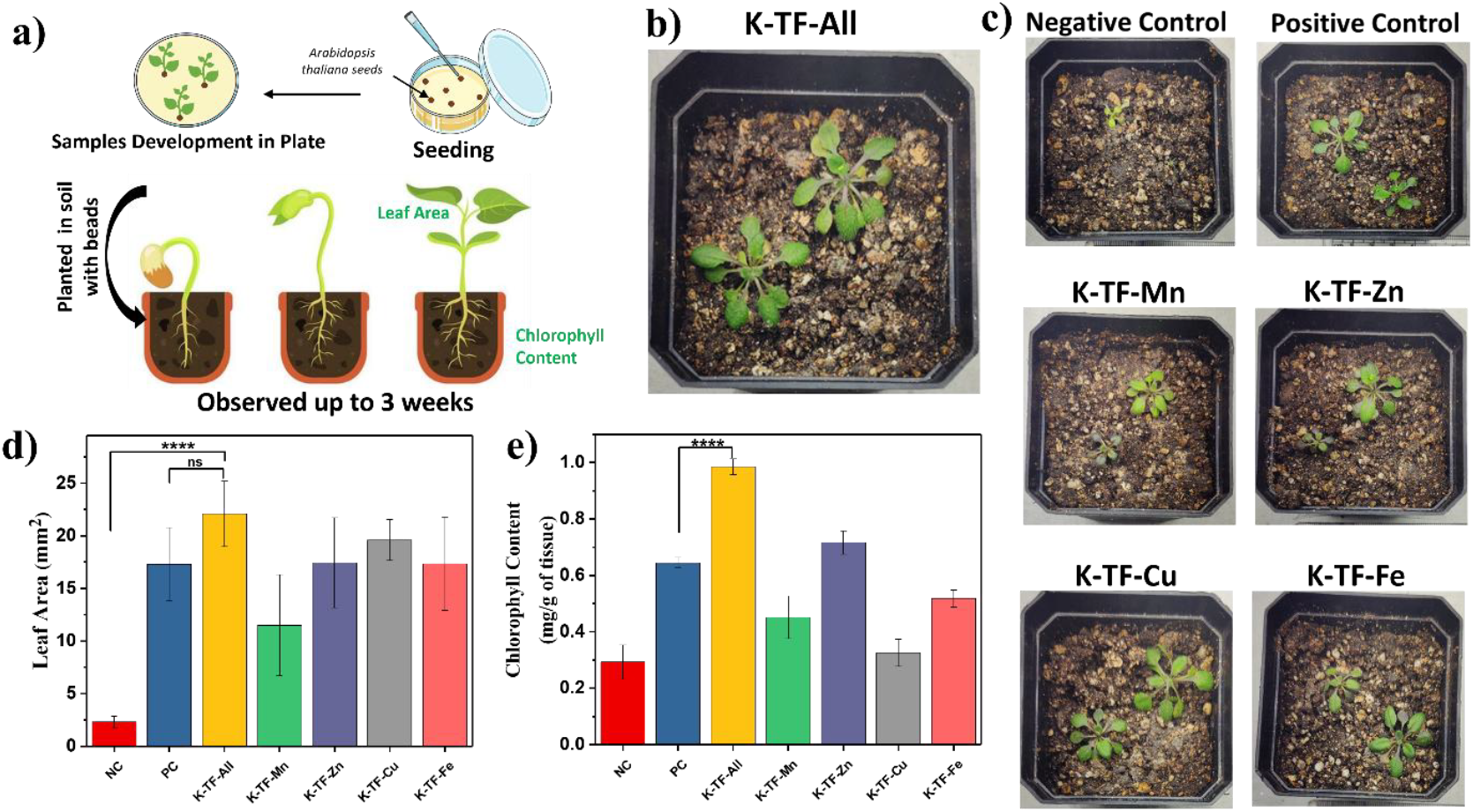
Evaluation of plant growth performance using *Arabidopsis thaliana*, showing (a) Schematic overview of the experimental procedure. (b) Photographs of plants treated with micronutrient-loaded beads after 21 days. (c) Comparative images of remaining treatment groups, including positive and negative controls, at day 21. (d) Leaf area analysis of plants subjected to various micronutrient treatments. (e) Chlorophyll content of plants treated with different micronutrient-loaded beads. Error bars = mean ± SD (n = 4).

**Figure 6.**
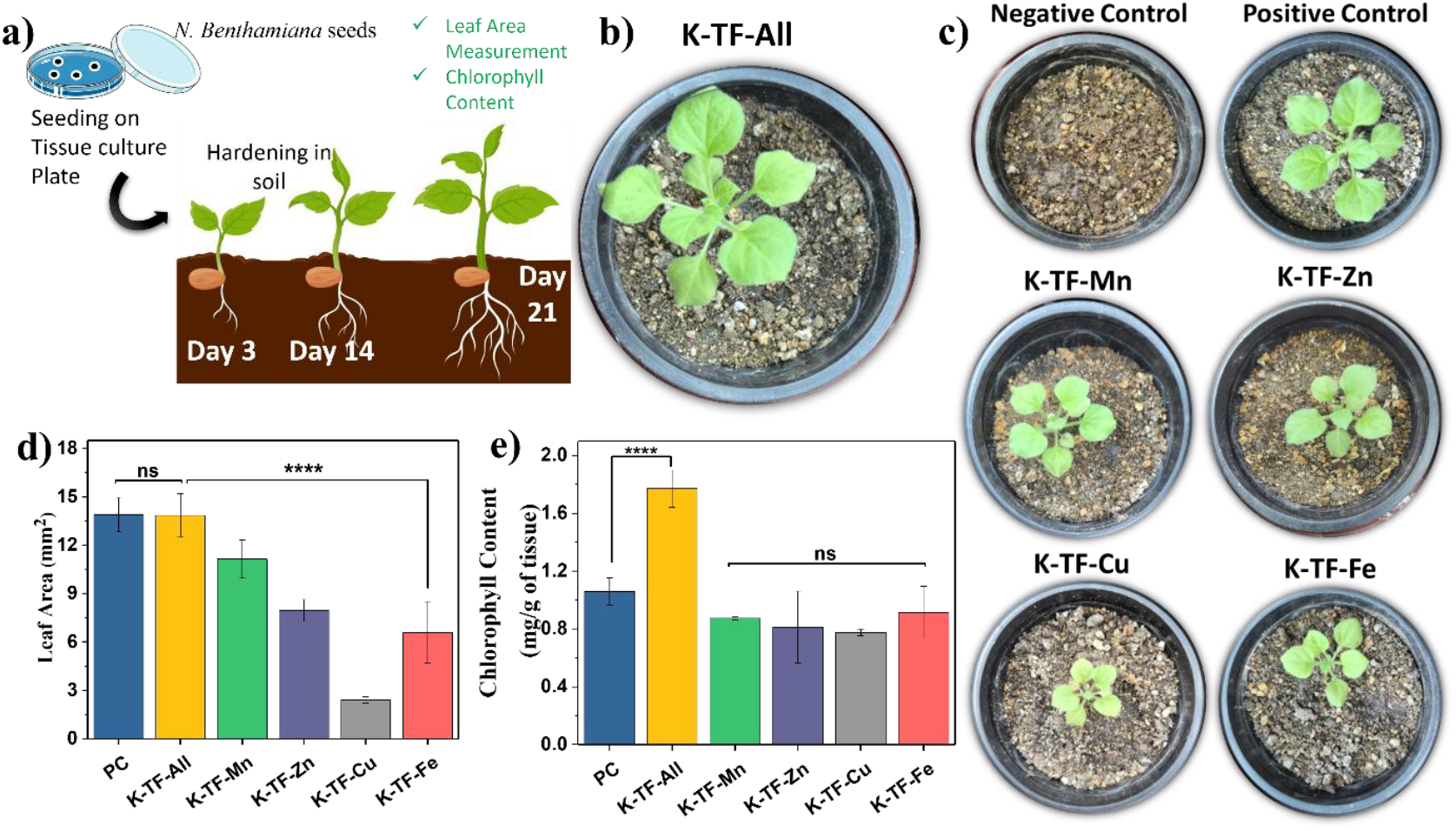
Assessment of plant growth response in *Nicotiana benthamiana*, illustrating (a) schematic representation of the experimental workflow. (b) Photographs of plants treated with micronutrient-loaded beads after 14 days. (c) Images of all remaining treatment groups, including positive and negative controls, at day 14. (d) Leaf area measurements of plants under various micronutrient treatments. (e) Chlorophyll content of plants treated with different micronutrient-loaded beads. Error bars = mean ± SD (n = 4).

#### Leaf area measurements

The Leaf area measurements revealed distinct variations across the treatments (**Figure 5b-d**). *Arabidopsis thaliana* treated with all micronutrient beads (K-TF-All) exhibited the largest leaf area (≈22 mm^2^), significantly greater than the negative control (NC) (≈2 mm^2^, ****p < 0.0001), indicating that the combined and balanced supply of micronutrients strongly promoted leaf expansion. Although the K-TF-All treatment was not significantly different from the positive control (PC) (≈18 mm^2^, ns), the numerical trend suggested a slight improvement over PC, reflecting the potential benefit of sustained micronutrient release from beads. Among the single-element treatments, Cu (K-TF-Cu), Fe (K-TF-Fe), and Zn (K-TF-Zn) beads supported intermediate leaf areas ranging from ≈17-20 mm^2^, comparable to PC, suggesting that these elements individually contribute to leaf growth but are less effective than the combined treatment. In contrast, Mn beads (K-TF-Mn) resulted in a considerably smaller leaf area (≈12 mm^2^), second only to the NC, highlighting the limited effect of Mn supplementation alone. Overall, these results indicate that while individual micronutrients induce moderate leaf expansion, the integrated delivery of all micronutrients via the microbeads produces the most pronounced growth response. Plants were monitored until the flowering stage, during which only the K-TF-All formulation and the positive control successfully reached flowering. Notably, plants treated with K-TF-All exhibited superior overall health compared to the positive control, which showed signs of drying, as evidenced in **Figure S2**. In *Nicotiana benthamiana* (**Figure 6 b-d**) the K-TF-All treatment produced the largest leaf area (≈14 mm^2^), which was comparable to the positive control and significantly greater than the other micronutrient treatments (**p < 0.0001**). All other micronutrient combinations resulted in smaller leaf areas, with K-TF-Cu bead treatment showing the smallest leaf area (≈2.5 mm^2^). Overall, these results indicate that integrated micronutrient delivery via K-TF-All is more effective than individual micronutrient supplementation in promoting leaf expansion in *N. benthamiana*.

#### Chlorophyll content analysis

*Arabidopsis thaliana* treated with all micronutrient beads accumulated the highest chlorophyll content, which was significantly greater than that of the positive control (p < 0.0001), underscoring the superior effectiveness of the bead-based delivery system (**Figure 5e**). While the PC provided a complete nutrient solution under optimal conditions, the K-TF-All treatment outperformed it, suggesting that the mode of nutrient delivery plays a critical role in enhancing plant physiology. The beads likely improved chlorophyll biosynthesis by providing a slow and pH-responsive release of micronutrients, ensuring a continuous and balanced supply to the plants throughout the growth period. In contrast, conventional supplementation in the PC may have led to a rapid availability of nutrients, followed by fluctuations or partial depletion over time. The controlled release from the beads would have minimized nutrient loss and potential antagonistic interactions, thereby improving uptake efficiency. This sustained micronutrient availability likely explains the higher chlorophyll accumulation observed in the K-TF-All treatment compared to PC. Together, these findings highlight that the synergistic effect of balanced micronutrient availability, combined with controlled release properties of the beads, provides a distinct advantage over traditional nutrient supplementation. A similar trend was observed in *Nicotiana benthamiana* (**Figure 6e**), where plants treated with all micronutrient beads exhibited the highest chlorophyll content (≈1.7mg/g of tissue), markedly exceeding that of the positive control as well as all single-element and combined treatments, which ranged from ≈0.7 to 1.0. Notably, all negative control plants failed to survive and therefore could not be included in chlorophyll analysis and leaf area measurement. Despite species-specific differences in absolute chlorophyll levels, the consistent enhancement observed with the All-beads treatment across both *Arabidopsis thaliana* and *Nicotiana benthamiana* highlights the robustness and cross-species applicability of the bead-based micronutrient delivery strategy.

## 4. Conclusion

In this study, we developed a biodegradable, pH-responsive microbead system based on κ-carrageenan and trans-ferulic acid for the controlled delivery of essential micronutrients (Mn^2+^, Zn^2+^, Cu^2+^, and Fe^3+^) under acidic soil environments. The fabricated microbeads exhibited desirable physicochemical properties, including uniform morphology, high encapsulation efficiency, and pronounced swelling and degradation under weakly acidic pH conditions simulating the rhizosphere. Nutrient release studies revealed a distinct pH-dependent behavior, with maximal cumulative release at pH 4, confirming the system’s responsiveness to rhizospheric acidification driven by root exudates. Spectroscopic analyses and antioxidant assays validated the structural stability and functional integrity of the carrageenan-phenolic matrix, while soil leaching experiments demonstrated the formulation’s environmental applicability and reduced nutrient losses. Importantly, plant bioassays with *Arabidopsis thaliana* and *Nicotiana benthamiana* confirmed the agronomic potential of the system, as evidenced by improved seedling vigor, enhanced leaf development, and the absence of phytotoxic effects.

Collectively, these findings establish the κ-carrageenan-trans-ferulic acid microbeads as a promising platform for targeted micronutrient delivery in acidic and nutrient-deficient soils, with potential to mitigate hidden hunger and promote sustainable plant nutrition. Future studies should focus on field-scale validation, long-term impacts on soil health and microbial ecology, quantification of nutrient bioavailability, and integration with macronutrients and beneficial microbes. Ultimately, this approach offers a foundation for the development of next-generation smart fertilizers tailored for precision agriculture and environmental resilience.

## Acknowledgments

MD thanks the Indian Institute of Technology Gandhinagar for the Seed Grant (IP/IITGN/BE/MD/2223/13). SS thanks DBT-RL (RES/DBTRL/BE/PO369/2223/0018) and

IITGN startup grant (Project IP/IITGN/BE/SS/2223/17). The authors are appreciative of the financial assistance provided by IIT Gandhinagar through research funding and Ph.D. fellowships. We also thank the Central Instrumentation Facility (CIF) for assistance with hydrogel characterization. The authors further acknowledge the use of AI-based tools solely for refining the language and structure of the manuscript, with no role in data interpretation or analysis.

## Data availability statement

All the data related to the study has already been provided.

## References

1. Sekar, N., & Ramasamy, R. P. (2015). Recent advances in photosynthetic energy conversion. Journal of Photochemistry and Photobiology C: Photochemistry Reviews, 22, 19–33. DOI: 10.1016/j.jphotochemrev.2014.09.004

2. Bhat, M. A., Mishra, A. K., Shah, S. N., Bhat, M. A., Jan, S., Rahman, S., … & Jan, A. T. (2024). Soil and mineral nutrients in plant health: A prospective study of iron and phosphorus in the growth and development of plants. Current Issues in Molecular Biology, 46(6), 5194–5222. DOI: 10.3390/cimb46060312

3. Ekele, J. U., Webster, R., Perez de Heredia, F., Lane, K. E., Fadel, A., & Symonds, R. C. (2025). Current impacts of elevated CO2 on crop nutritional quality: a review using wheat as a case study. Stress Biology, 5(1), 34. DOI: 10.1007/s44154-025-00217-w

4. Shukla, A. K., Behera, S. K., Prakash, C., Tripathi, A., Patra, A. K., Dwivedi, B. S., … & Singh, A. K. (2021). Deficiency of phyto-available sulphur, zinc, boron, iron, copper and manganese in soils of India. Scientific reports, 11(1), 19760. DOI: 10.1038/s41598-021-99040-2

5. Solangi, F., Zhu, X., Cao, W., Dai, X., Solangi, K. A., Zhou, G., & Alwasel, Y. A. (2024). Nutrient uptake potential of nonleguminous species and its interaction with soil characteristics and enzyme activities in the agro-ecosystem. ACS omega, 9(12), 13860–13871. DOI: 10.1021/acsomega.3c08794

6. Oburger, E., & Schmidt, H. (2016). New methods to unravel rhizosphere processes. Trends in plant science, 21(3), 243–255. DOI: 10.1016/j.tplants.2015.12.005

7. Ma, W., Tang, S., Dengzeng, Z., Zhang, D., Zhang, T., & Ma, X. (2022). Root exudates contribute to belowground ecosystem hotspots: A review. Frontiers in Microbiology, 13, 937940. DOI: 10.3389/fmicb.2022.937940

8. Sica, P., Kopp, C., Müller-Stöver, D. S., & Magid, J. (2023). Acidification and alkalinization pretreatments of biowastes and their effect on P solubility and dynamics when placed in soil. Journal of Environmental Management, 333, 117447. DOI: 10.1016/j.jenvman.2023.117447

9. Kaźmierczak, M., Błońska, E., Kempf, M., Zarek, M., & Lasota, J. (2025). Rhizosphere effect: microbial and enzymatic dynamics in the rhizosphere of various shrub species. Plant and Soil, 511(1), 245–262. DOI: 10.1007/s11104-024-06981-4

10. Huang, K., Li, M., Li, R., Rasul, F., Shahzad, S., Wu, C., … & Aamer, M. (2023). Soil acidification and salinity: the importance of biochar application to agricultural soils. Frontiers in Plant Science, 14, 1206820. DOI: 10.3389/fpls.2023.1206820

11. Hasegawa, H., Nozawa, A., Papry, R. I., Maki, T., Miki, O., & Rahman, M. A. (2018). Effect of biodegradable chelating ligands on Fe uptake in and growth of marine microalgae. Journal of Applied Phycology, 30(4), 2215–2225. DOI: 10.1007/s10811-018-1462-x

12. Mim, J. J., Rahman, S. M., Khan, F., Paul, D., Sikder, S., Das, H. P., … & Hossain, N. (2025). Towards smart agriculture through nano-fertilizer-A review. Materials Today Sustainability, 30, 101100. DOI: 10.1016/j.mtsust.2025.101100

13. Zhang, D. X., Du, J., Wang, R., Luo, J., Jing, T. F., Li, B. X., … & Hou, Y. (2021). Core/shell dual-responsive nanocarriers via iron-mineralized electrostatic self-assembly for precise pesticide delivery. Advanced Functional Materials, 31(34), 2102027. DOI: 10.1002/adfm.202102027

14. Hou, X., Pan, Y., Xiao, H., & Liu, J. (2019). Controlled release of agrochemicals using pH and redox dual-responsive cellulose nanogels. Journal of agricultural and food chemistry, 67(24), 6700–6707. DOI: 10.1021/acs.jafc.9b00536

15. Antonopoulou, I., Sapountzaki, E., Rova, U., & Christakopoulos, P. (2022). Ferulic acid from plant biomass: A phytochemical with promising antiviral properties. Frontiers in nutrition, 8, 777576. DOI: 10.3389/fnut.2021.777576

16. Gupta, R. K., Rajan, D., Meena, D., & Srivastav, P. P. (2025). Ferulic Acid as a Sustainable and Green Crosslinker for Biopolymer-Based Food Packaging Film. Macromolecular Chemistry and Physics, 226(10), 2400441. DOI: 10.1002/macp.202400441

17. Shahabadi, N., Karampour, F., Fatahi, N., & Zendehcheshm, S. (2022). Synthesis, characterization, in vitro cytotoxicity and DNA interaction studies of antioxidant ferulic acid loaded on γ-Fe2O3@ SiO2 nanoparticles. Nucleosides, Nucleotides & Nucleic Acids, 41(10), 994–1011. DOI: 10.1080/15257770.2022.2094409

18. Mancuso, C., & Santangelo, R. (2014). Ferulic acid: pharmacological and toxicological aspects. Food and Chemical Toxicology, 65, 185–195. DOI: 10.1016/j.fct.2013.12.024

19. Zia, K. M., Tabasum, S., Nasif, M., Sultan, N., Aslam, N., Noreen, A., & Zuber, M. (2017). A review on synthesis, properties and applications of natural polymer based carrageenan blends and composites. International journal of biological macromolecules, 96, 282–301. DOI: 10.1016/j.ijbiomac.2016.11.095

20. Kadota, K., Nogami, S., Uchiyama, H., & Tozuka, Y. (2020). Controlled release behavior of curcumin from kappa-carrageenan gels with flexible texture by the addition of metal chlorides. Food Hydrocolloids, 101, 105564. DOI: 10.1016/j.foodhyd.2019.105564

21. Farooq, A., Farooq, A., Jabeen, S., Islam, A., Gull, N., Khan, R. U., … & Bilal, M. (2022). Designing Kappa-carrageenan/guar gum/polyvinyl alcohol-based pH-responsive silane-crosslinked hydrogels for controlled release of cephradine. Journal of Drug Delivery Science and Technology, 67, 102969. DOI: 10.1016/j.jddst.2021.102969

22. Singh, S., & Pal, K. (2024). Polyphenol modified CuO nanorods capped by kappa-carrageenan for controlled paclitaxel release in furnishing targeted chemotherapy in breast carcinoma cells. International Journal of Biological Macromolecules, 255, 127893. DOI: 10.1016/j.ijbiomac.2023.127893

23. Vithalani, H., Dave, H., Singh, H., Sharma, D., Navale, A., & Dhanka, M. (2025). Mechanically robust, mouldable, dynamically crosslinked hydrogel flap with multiple functionalities for accelerated deep skin wound healing. Biomaterials Advances, 169, 214195. DOI: 10.1016/j.bioadv.2025.214195

24. Chugh, P., & Sharma, P. (2021). Effect of heat stress on the relationship between SPAD and chlorophyll content in Indian mustard geno types. Current Journal of Applied Science and Technology, 40(15), 38–17. DOI: 10.9734/cjast/2021/v40i1531412.

