## Supplemental file for "Engineering pH-Responsive trans-Ferulic Acid/κ-Carrageenan Beads for On-Demand Micronutrient Delivery in Plants"

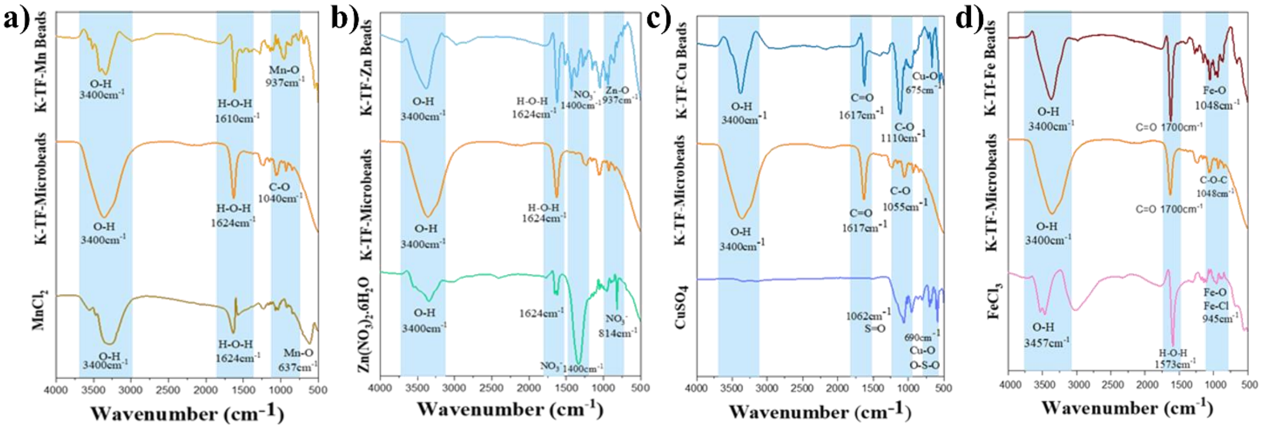


**Figure S1: FTIR Spectra of different bead formulations along with their controls (a) K-TF-Mn, (b) K-TF-Zn, (c) K-TF-Cu, (d) K-TF-Fe**

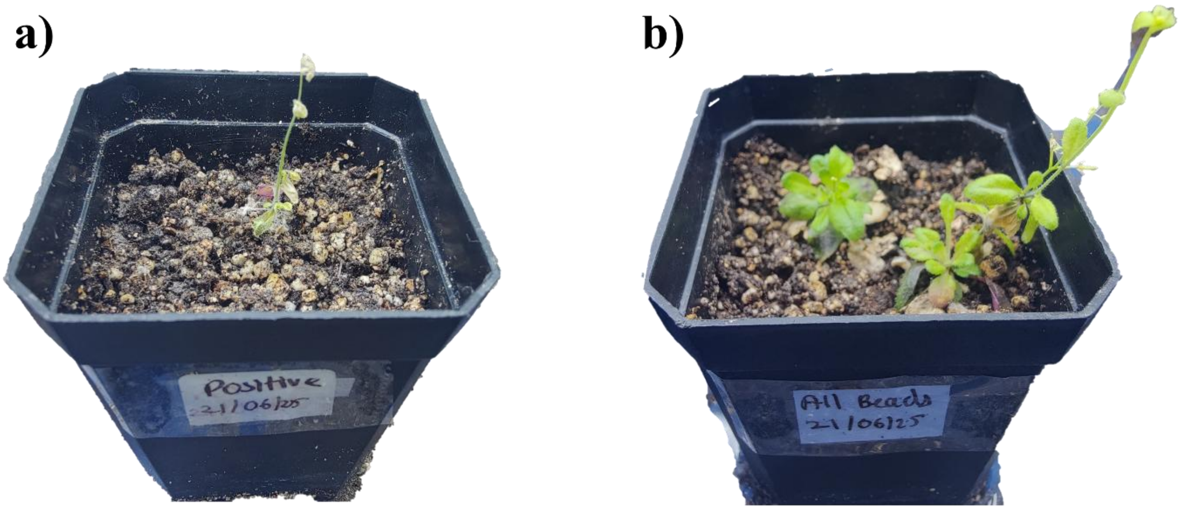


**Figure S2: Flowering stage in *Arabidopsis thaliana* (a) Positive control, (b) K-TF-All**
